# Efficacy of Topical Strategies to Prevent the Onset of Root Caries Lesions, *In vitro*

**DOI:** 10.1101/2024.02.19.581042

**Authors:** Ricardo Valenzuela-Ramos, Soraya León, Rodrigo A. Giacaman

## Abstract

Preventive fluoride-based topical products, such as silver diamine fluoride (SDF) and others with chlorhexidine have been widely used as a strategy for preventing dental caries lesions, at all ages. Whether they are effective against root caries is less clear.

**Aim:** To evaluate the efficacy of various dental products in preventing root caries lesions, using a biological model of root caries.

**Methodology:** An experimental *in vitro* study was devised using a validated model with biofilms of *Streptococcus mutans* formed on bovine root dentin slabs. Slabs were divided into 5 experimental groups: (1) 1% chlorhexidine gel, (2) 22,600 ppm fluoride varnish, (3) 5000 ppm fluoride toothpaste, (4) 38% SDF and (5) negative control (0.9% NaCl). After the experimental phases, biofilms were detached from the dentin slabs to assess biomass and the number of viable cells (CFU). Demineralization of dentin slabs was evaluated by laser fluorescence and surface Knoop microhardness loss.

**Results:** Dentin slabs treated with SDF did not significantly differ in surface microhardness loss compared to those treated with 1% chlorhexidine gel (p>0.05). However, both materials showed significantly lower surface demineralization than the rest of the groups. SDF had a higher impact on the properties of *Streptococcus mutans* biofilms.

**Conclusion:** SDF seems more effective than other materials in preventing root caries lesions in a highly cariogenic environment.

## Introduction

Untreated dental caries is the most common oral condition among adults worldwide, with its incidence increasing after the age of 60, primarily due to the rise of root caries in the elderly population [1]. The prevalence of root caries in individuals aged 60 and older is estimated to range from 8% to 74% [2]. The widespread prevalence of root caries in older individuals leads to a peak in untreated dental caries around the age of 70 in the global adult population, resulting in a significant economic burden on society and governments [3].

Furthermore, untreated caries has a negative impact on the quality of life of older individuals, especially due to the experience of pain, which can lead to psychological and physical discomfort, and even disability [4]. Residents in nursing homes are at a higher risk of developing dental caries, even more so than individuals of the same age who are not institutionalized [5]. Even measures such as providing access to preventive dental caries treatment for institutionalized older adults, including fluoride varnish applications up to four times a year, are deemed insufficient [6]. Therefore, having effective methods for preventing root caries becomes a crucial factor in reducing its incidence and improving the quality of life for these patients.

Fluoride therapy has been the most effective strategy in preventing dental caries lesions [7, 8]. Fluoride-releasing agents are available both as preventive methods and as early non-invasive treatment of caries lesions [9–15], the evidence supports that the application of a high concentration fluoride (F) is more effective than the use of a standard fluoride toothpaste [9]. Regular tooth brushing with fluoride dentifrices is the most rational way to administer topical F. Toothpaste with 5000 ppm F applied twice daily has been reported to be more effective than dentifrice with 1450 ppm F in stopping root caries lesions in both dependent individuals applied by nursing staff [16], and applied autonomously by patients [9]. Although it is an effective method, its application requires the cooperation of the patient or the caregiver. Therefore, it cannot be the best first-choice preventive treatment in frail, cognitively impaired, or institutionalized patients, where correct oral hygiene is diminished by its biopsychosocial characteristics.

While much of the current literature concerning fluoride varnishes pertains to their role in preventing coronal caries in children and young adults, a study has shed light on their efficacy in a different demographic. Specifically, in institutionalized older adults, the application of 5% NaF varnish every 3 months resulted in a remarkable 64% reduction in root caries incidence during the 36-month follow-up period. Additionally, this study found both 5% NaF varnish and 38% silver diamine fluoride (SDF) solution to be equally effective in preventing root caries, with no statistically significant differences observed between the two agents [17].

During the last decade, silver diamine fluoride (SDF) has had a prominent role as an effective preventive method of dental caries, both in enamel caries [18–22], as on root lesions [23–29]. SDF is an alkaline topical solution that contains fluoride and silver [30]. Its benefits include having the highest fluoride concentration among all fluoride products used in dentistry [31], reducing the growth of cariogenic bacteria [32–35] and promote the remineralization of the inorganic content of enamel and dentin. With this, SDF prevents the degradation of collagen in dentin by inhibiting the activity of collagenases and cysteine-cathepsins [31]. Likewise, it has been reported that dentin treated with SDF remineralizes and releases fluoride for up to 48 hours [20]. Given the above, it has been described as the most effective, efficient and safe agent to control caries in both primary and permanent dentition compared to any other fluoride agent [36], being an effective measure, and low cost. Despite the numerous benefits, the unesthetic black staining of the carious tissue hampers the broader acceptance of SDF, mainly by dentists, but not patients [37]. A recent systematic review showed that 5% NaF varnish and 38% SDF were the most effective agents in preventing root caries, with no significant differences between them [38]. Another non-invasive product for topical use as a preventive method for root caries is 1% chlorhexidine gel (CHX). Evidence shows that CHX can inhibit the growth of cariogenic bacteria on the root surface [39–43], specifically on S. mutans [44], reducing root caries lesions [45, 46], and even as a tertiary preventive method, showing favorable results [45, 47, 48].

The existing evidence is insufficient to demonstrate which of the available agents is more efficient to counteract the cariogenic action of sucrose in healthy root tissues exposed to this etiological factor of dental caries. The present study compared the anticaries activity of different materials for topical use in preventing root caries. The selection of topical agents was based on their ease of application, particularly in populations with compromised functional capacities. This consideration is crucial as it ensures that maintaining functional oral health remains feasible even in circumstances where regular maintenance may be challenging.

## Materials and methods

A previously validated caries model with biofilms of the cariogenic *Streptococcus mutans* (*S. mutans*) UA159 was used [49]. This model allows assessing the efficacy of various dental materials in the prevention of root caries lesions, in a biological model of root caries upon several cariogenic challenges. The dentin slabs were treated with 5 types of root caries preventive topical materials (5 experimental groups (n=3): (1) 1% chlorhexidine gel, (2) 22,600 ppm fluoride varnish, (3) 5000ppm fluoride toothpaste, (4) 38% SDF and (5) negative control (0.9% NaCl)), followed by biofilm formation of *S. mutans* on the treated bovine dentin slabs. For 4 days, biofilms/slabs were exposed three times per day to a solution of 10% sucrose (w/v) for 5 min to create cariogenic conditions. Culture medium was replenished twice per day, and the pH of the spent medium was measured to evaluate acidogenicity elicited by the biofilm. After the 4 days of the experimental phase, biofilms were recovered to assess total biomass, viable bacteria, and biofilm soluble proteins. Dentin demineralization was estimated by surface Knoop microhardness (SH) loss and laser fluorescence readings (LF). Two independent experiments were carried out, each in triplicate (n = 6).

### Dental dentin samples

Bovine incisors were cleaned with a 5% NaOCl solution and stored in 0.9% NaCl (w/v) until use, avoiding storage times longer than thirty days. Thirty dentin slabs of 4 x 4 x 1 mm were obtained and polished with 500 and 1,200 grit sandpaper and finished using Soflex polishing discs (3M, St. Paul, MN, USA). To obtain an initial value for Knoop surface microhardness (SH_i_), three indentations were performed with a microindenter with a microhardness tester (402 MVD, Wolpert Wilson Instruments, USA) at 10 g for 5 sec. Mean baseline SHi of all the slabs was 60,81 ± 8,07 and only those with SH_i_ within a 10% variation were included. Thus, bias from including slabs with too different SHi was avoided. LF records for caries detection were performed with a DIAGNOdent pencil (KaVo, Biberach, Germany), One device and the same tip were used at each center throughout each measurement. All sites were measured three times on each occasion, and the mean of the values was recorded. Slabs were sterilized with ethylene oxide [50].

### Treatments and *Streptococcus mutans* biofilms

The dentin slabs were distributed into 5 groups (n=3), and were treated with root caries preventive topical materials, following the manufacturer’s instructions. The experimental groups were: (1) 1% chlorhexidine gel (Cervitec P, Ivoclar Vivadent, Schaan Liechtenstein), (2) 22,600 ppm fluoride varnish (Clinpro Varnish 3M, Saint Paul, MN, USA), (3) 5000 ppm fluoride dentifrice (Caristop 5000, Maver, Santiago, Chile), (4) 38% silver diamine fluoride (FAGAMIN, Tedequim, Cordova, Argentina) and (5) negative control (0.9% NaCl).

An acquired pellicle-like structure was created on the dentin root surface treated with caries-preventive materials, with the purpose of emulating the situation in vivo and also to facilitate bacterial adhesion. Slabs were immersed in ultrafiltered (0,22 μm) pooled human saliva for 30 min with a protease inhibitor cocktail (Koo et al. 2003). Initially measured, saliva-coated and sterile slabs were placed into a 24-well cell culture plate suspended by sterile wire.

*Streptococcus mutans* biofilms were formed on the dentin slabs, following a previously reported protocol [49]. Dentin slabs were inoculated with *S. mutans* cultures (OD 0,8 at 600 nm) and 1% sucrose-containing medium to form the adherent biofilm [51] and incubated for 8 h. Biofilms were allowed to mature on the slabs in BHI medium supplemented with 0·1 m mol l−1 glucose for 24 h.

For 4 days, biofilms/slabs were exposed three times per day to a solution of 10% sucrose (w/v) for 5 min to create cariogenic conditions. Immediately after cariogenic challenge with 10% sucrose, blocks and biofilms were washed with 0.9% NaCl three times and replated on the plate.

### Biofilm acidogenicity

To verify acidogenicity of the biofilm, culture medium pH was measured in the wells. A pH meter (Orion Star, Thermo Scientific) was used to register the pH. Spent culture medium generated by the biofilm was read twice per day, before each medium change [49].

### Biofilm analysis

To determine the microbial composition, biofilm-containing slabs were transferred to 1 mL 0.9% NaCl. The biofilms were detached from the slabs by sonication at 7W for 30 seconds and the slabs stored for further analysis. The biofilm suspension was aliquoted to evaluate the following biofilm properties: biomass, bacterial counting, and biofilm soluble proteins, briefly described as follows:

### Biomass

To estimate biomass, 200 µL from the biofilm suspension were transferred to a pre-weighted tube and incubated with 100% ethanol at -20°C for 15 min, centrifuged (10 min at 5000 g and 4°C) and the resulting pellet was washed with 500 µL of 75% ethanol, centrifuged again and dried for 24 h in a desiccator. Biomass was calculated by subtracting the final weight to the initial weight of the empty tube. Biomass dry weight of the biofilm was expressed as mg per mL of biofilm suspension [51].

### Viable bacteria in the biofilm

To obtain the number of viable colonies, a 100 µL aliquot of the biofilm suspension was serially diluted up to 1: 14,000 (v/v) in 0.9% NaCl at pH 7.0 and each dilution were drop-plated on BHI agar plates in duplicate. Plates were incubated anaerobically for 24 h at 37°C, and colonies were enumerated from the dilution that allowed visualization of isolated colonies. Counting was corrected by the dilution factor and expressed as CFU mg−1 of biofilm dry weight [52].

### Biofilm soluble proteins

Fifty µL aliquots of the biofilm suspension were treated with 2 mol−1 NaOH and incubated for 15 min at 100°C. Samples were centrifuged at 10,000 g at 4°C for 10 min, and the supernatant was used to assess the total protein concentration of the biofilm by the Lowry’s method (Lowry et al. 1951), using Bradford’s reagent (Sigma, USA). Results were expressed as µg of protein per mg of biomass.

### Enamel demineralization

Using the recovered slabs, SH loss was used as a surrogate indicator of enamel demineralization [53]. SH has been extensively used as a reliable methodology to evaluate demineralization [54]. Briefly, final indentations were performed 100 μm away from those performed before the experimental phase to obtain a final SH reading (SH_f_ in Kg/mm^2^) in a triplicate row. Mean values from the initial and final measurements were used to obtain the percentage of SH loss (%SHL) calculated by the formula: (SH_i_ – SH_f_)*100/SH_i_.

### Laser fluorescence readings

LF records for caries-like lesion detection were performed using the DIAGNOdent pencil (KaVo, Biberach, Germany) at baseline and at the end of the experiment. One device and the same tip were used at the same spot before and after the experiments, using the center portion of the slab. All sites were measured three times on each occasion, and the mean values were recorded. The instrument was calibrated between each slab measurement using a ceramic standard according to the manufacturer’s instructions.

### Statistical analysis

Parametric distribution of the data was verified by the Kolmogorov–Smirnov test. Dependent quantitative variables were analyzed by ANOVA test. When differences were found, a post-hoc Bonferroni test followed to test for differences between pairs of treatment groups. The SPSS 29.0 statistical software, (IBM Corporation, Somers, NY) was used to analyze the data, setting a 95% confidence level.

## Results

The sucrose-induced demineralization as a cariogenic challenge to the biofilm revealed that dentin slabs treated with SDF resulted in significantly lower demineralization compared to all study groups (p<0.05). Dentin slabs treated with CHX, prior to the sucrose challenge, only showed significantly lower demineralization, when compared to the 5000 ppm F dentifrice and to the control group (Figure 1). Fluoride varnish only exhibited lower demineralization compared to the control group (p<0.05) (Figure 1A). When estimated through LF, demineralization was significantly lower for SDF compared to the other groups, but without differences with CHX (p>0.05). Consistent with the %SHL assay, the CHX group displayed lower values than the other groups, but with non-significant differences (p>0.05) (Figure 1B).

**Figure 1:**
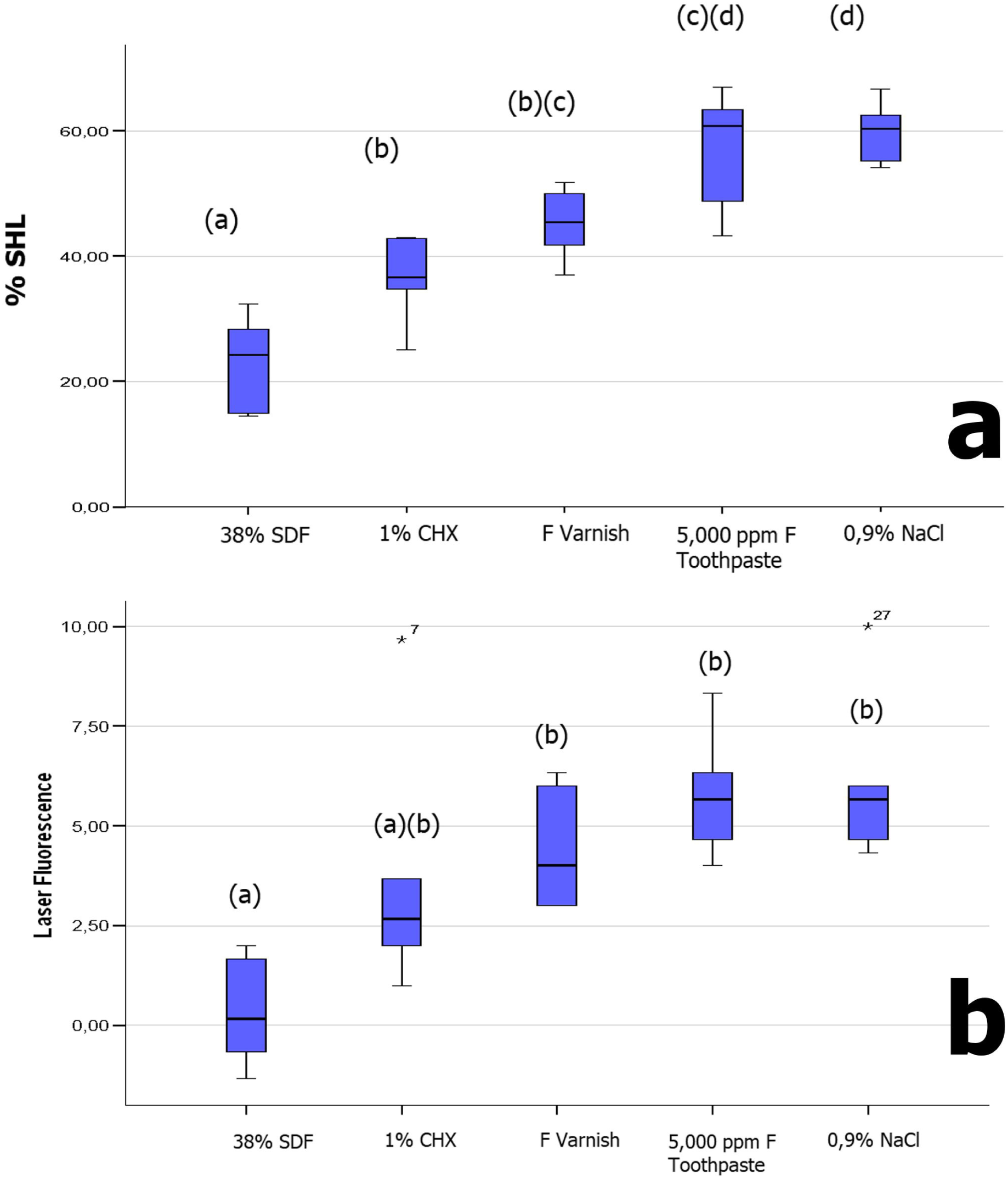
Demineralization of treated dentin, in response to sucrose exposure to the *S. mutans* biofilms. (a) Percentage of surface hardness loss (%SHL). Surface microhardness (SH) was measured before and after the experiment to evaluate the %SHL. (b) Demineralization estimated through laser fluorescence (LF) (DIAGNOdent pen) at baseline and at the end of the experiment. Box plots represent values from 2 independent experiments in triplicate (n=6). Different letters indicate statistically significant differences (p<0.05).

Acidogenicity induced by the biofilms formed on the slabs was assessed through pH measurements of the medium, twice daily throughout the experiment. SDF-treated biofilms showed higher pH than the rest of the groups throughout the experiment (p<0.05), except for CHX that did not have differences with SDF during the initial 48 h (p>0.05). However, from 56 h, only SDF displayed a significantly higher pH than the rest of the groups (p<0.05) (Figure 2).

**Figure 2:**
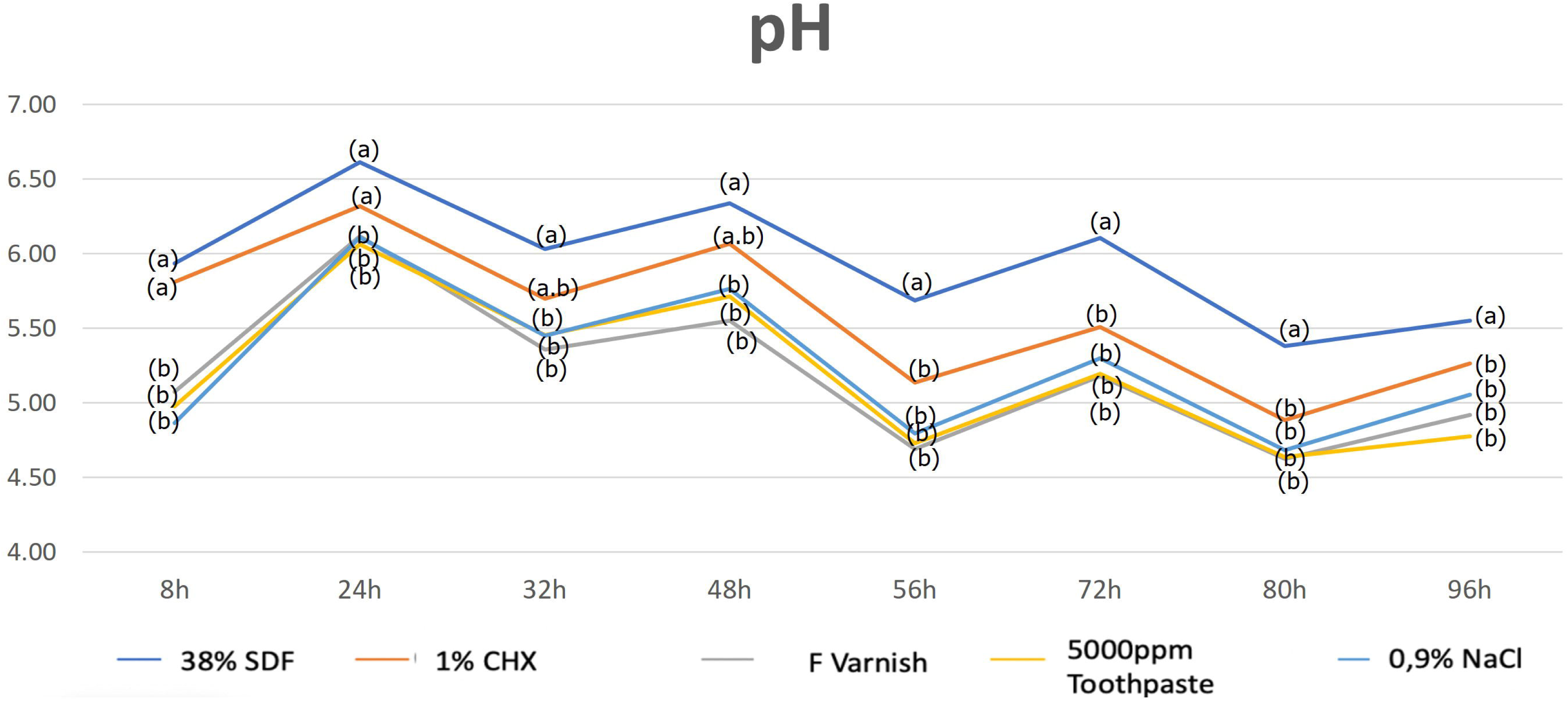
Acidogenicity of *S. mutans* biofilms exposed to different treatments. Spent medium was measured for pH at indicated times. Each time-point in the plot represents the mean of 2 independent experiments in triplicate slabs. Medium changes are represented by the arrows. Different letters at each time point represent statistical significant differences (p<0.05).

SDF pretreated biofilms on dentin slabs exhibited significantly (p<0.05) lower counts of viable cells than all the rest of the experimental groups. CHX, Fluoride varnish and 5,000 ppm F groups did not disclose differences among them (p>0.05), with CHX resulting in less CFU when compared to the control with 0.9% NaCl (Figure 3a). Similarly, only the SDF group showed significantly lower amounts of biomass compared to all other study groups (p<0.05), except for CHX that showed no differences with SDF. Fluoride varnish and 5,000 ppm F failed to show differences with the caries-negative control (p>0.05) (Figure 3b). Conversely, protein content in the biofilms did not reveal differences in the treatment groups (p>0.05), all of them having lower values than the negative control (p<0.05). (Figure 3c).

**Figure 3:**
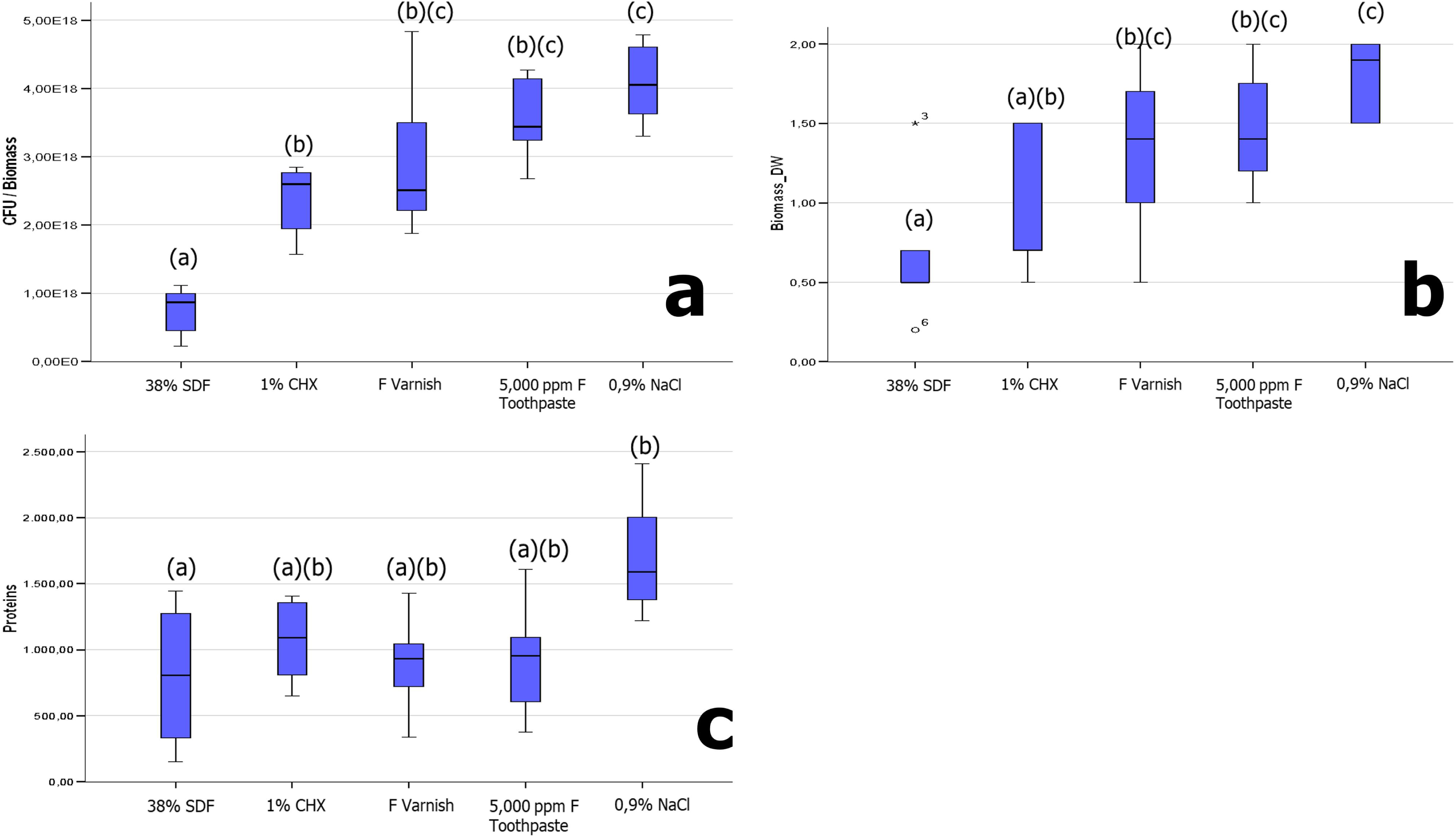
(a) number of colony-forming units (CFU) of *S. mutans* in the biofilms of the pretreated slabs. (b) Biomass by the dry-weight method (c) Total protein concentration by the Lowry’s method. Box-plots represent values from 2 independent experiments in triplicate (n=6). Different letters indicate statistically significant differences (p<0.05).

## Discussion

This study tested different preventive topical materials, used generally for caries prevention, in a root caries experimental model with biofilms. The main finding was that SDF reduced root caries-like lesion formation and inhibited biofilm metabolic activity. CHX gel also exhibited a caries-inhibitory effect, more restricted than SDF, but more pronounced than the other experimental groups; 5,000 ppm F dentifrice and fluoride varnish. These findings are in line with a systematic review that reported that 1% chlorhexidine gel and SDF appear to be more effective in preventing root caries, when compared to conventional fluoride toothpaste [55]. Consistently with our results, a meta-analysis revealed that both SDF and CHX are clinically effective in reducing new root caries lesions compared to placebos, but with SDF showing a better effect than chlorhexidine [56]. SDF has shown higher preventive fractions, when compared to other topical applications like NaF varnish [21, 22, 57], even when tested in dependent or institutionalized elderly individuals [24]. However, other authors have not found significant differences between the use of fluoride varnish and SDF [58], not even when adding SDF to fluoride varnish in institutionalized elderly individuals compared to using fluoride varnish alone [59].

Untreated dental caries lesions represent the most common oral condition in adults, and their incidence significantly increases after the age of 60, primarily due to the rising occurrence of root caries [1]. The prevention of these lesions involves maintaining proper oral hygiene, adhering to a healthy diet, and scheduling regular dental visits [60]. However, this is not always feasible, especially in frail elderly patients who may have difficulties in maintaining adequate oral hygiene due to underlying medical conditions. Therefore, employing local therapies capable of rendering the root surface more resistant to acid attacks and dental biofilm activity emerges as an enticing objective. The rationale behind this study was to identify a topical application of a preventive agent capable of fortifying tooth structure while simultaneously addressing cariogenic biofilm activity. Hence, the agents were administered prior to biofilm formation, simulating a clinical scenario where either the dentist or the patient applies the agent followed by biofilm formation. Our model aimed to simulate repeated exposure to sucrose, thus creating a highly cariogenic environment akin to that encountered in elderly individuals with limited biofilm removal abilities. This is an expanding area in dentistry and, indeed, recent reports indicate efforts towards testing new technologies aiming similar goals [61].

It was interesting to verify that the standard of care in preventing root caries lesions [29, 62]; the 5,000 ppm F dentifrice, showed lower capacities to control root caries than SDF or 1% CHX. The use of a dentifrice in this model, however, cannot compete with the topical agents, as the former needs to be applied daily and more than once daily to achieve optimal results, as we have demonstrated before [9]. There is a technical limitation in applying the 5,000 ppm F dentifrice in this model, for the biofilm would need to be removed and altered, which would create different experimental conditions for this group.

Consistent with these findings, the evidence has reported that 1% CHX gel significantly reduces root caries lesions [45, 46]. Moreover, it has even been recommended for root caries in patients requiring special care [44], such as institutionalized elderly people, as it can control established root lesions and reduce the incidence of new lesions [45]. However, the evidence remains unclear, as most research has been carried out in coronal caries on enamel. Furthermore and unlike SDF, studies have reported that CHX varnishes have no effect on the biofilm compared to fluoride varnish [43] and have no significant preventive effect on root caries compared to other types of fluoride [48, 63]. In the same line, a previous *in vitro* study focusing on *Lactobacillus rhamnosus*, showed that neither CHX rinses nor varnishes had a caries-protective effect on root caries [64]. An important observation was that dentin slabs treated with SDF or CHX gel not only exhibited reduced demineralization, but also showed a significantly lower number of CFU of *S. mutans*, suggesting a microbiological effect, even when the application is performed before the biofilm formation. The lower number of CFU of *S. mutans* in the biofilms of the slabs treated with SDF can be explained by the high concentration of ionized silver in SDF, which has a bactericidal effect on cariogenic microorganisms, specifically *S. mutans* [31, 65]. *In vitro* studies using biofilm models have shown that silver interacts with bacterial cell membranes and their enzymes, inhibiting bacterial growth [32, 66]. It is known that SDF, by reducing biofilm growth, increases the pH of the biofilm [67], which hinders collagen degradation [68]. It is interesting to note that the application of SDF in this study occurred prior to the formation of biofilms, therefore, the alkaline pH could act as a buffer for the culture medium, influencing the colonization of *S. mutans*. It would be interesting to separately evaluate the effect of pH on biofilms, and the antimicrobial properties of the silver ion of SDF in further studies. Likewise, 1% CHX gel also exhibited a decrease in the number of CFU of *S. mutans*, possibly due to the broad-spectrum antimicrobial properties of CHX [69]. Furthermore, CHX appears to exert a persistent antimicrobial activity against *S. mutans* when used in form of a gel [44].

Products containing fluoride offer varying levels of protection for root caries against demineralization. Based on our findings, it appears that traditional methods such as fluoride varnish and 5,000 ppm F are not sufficient in patients exposed to a highly cariogenic environment. Indeed, it has been reported that in institutionalized individuals or in those with severe dependency, conventional methods like fluoride varnish [70], or traditional toothpaste are inadequate for maintaining good oral health [27]. This study demonstrates an improved effect of SDF in preventing root caries, when modeled using an *in vitro* approach, consistent with a recent systematic review [15]. Besides an anticaries effect, SDF can be considered an effective cariostatic agent, easy to apply, affordable, and safe, making it suitable for inclusion in minimally invasive dentistry strategies [71]. Concerning the primary drawback of SDF therapy, the staining that primarily manifests in active dental caries poses minimal concern in healthy enamel and root dentin, as it is typically minor and can be effectively removed through polishing [72]. On the other hand, CHX gel appears to be also effective, but not better than SDF, especially regarding its influence on the biofilm. It is important to mention that SDF is more affordable than CHX gel, which could be a major issue, when thinking of these measures as a public health policy.

We acknowledge the limitations implicit in this study. The results should be carefully interpreted as they stem from an *in vitro* study employing a single-species biofilm. Dental caries, including root caries is a non-communicable disease mediated by biological factors, mainly sugars consumption, but also by a myriad of other systemic and social variables, intrinsically rooted in human behaviors [73]. Hence, applying single measures may not be as efficient as when including caries in a more holistic and interdisciplinary health care strategy [73]. Therefore, these findings must be treated as proof-of-principle leading to further clinical trials.

## Conclusion

The findings of this study suggest that SDF demonstrates notable efficacy in preventing root caries lesions, especially when used in situations characterized by high cariogenic activity. Similarly, the application of 1% Chlorhexidine (CHX) gel appears to yield results comparable to SDF, showcasing superiority over conventional fluoride varnish and dentifrice containing 5,000 ppm fluoride. However, to substantiate these observations and ascertain the generalizability of these treatments, further clinical trials are warranted, particularly focusing on vulnerable populations such as frail or dependent elderly individuals.

## Acknowledgments

The authors thank Ms. Natalia García-Manríquez for her technical assistance in the experiments. This study was presented as part of the graduation requirements of RV in the Master’s graduate program in Geriatric Dentistry of the University of Talca. The authors declare no potential interest with the products cited in this study.

## Study approval statement

This study protocol did not require ethical approval.

## Conflict of Interest Statement

The authors have no conflicts of interest to declare.

## Funding Sources

The study was funded by the Chilean Government Grant Fondecyt Regular 1210188 to RAG.

## Author Contributions

RV designed and conducted the experiments, acquired and analyzed the data, and drafted the first manuscript. RAG conceived the idea, designed the experimental setting and wrote the final version of the manuscript. SL contributed to the data analysis and discussion and critically revised the manuscript. All authors revised and approved the final version of the article.

## Data Availability Statement

All data generated or analyzed during this study are included in this article. Further enquiries can be directed to the corresponding author at.

## Notes

### Competing Interest Statement

The authors have declared no competing interest.

